# Lesion-Gradient Mapping in Semantic Aphasia: Comparisons of Observed and Simulated Effects of Stroke on Connectivity Gradients

**DOI:** 10.64898/2026.06.12.731654

**Authors:** Ramya Balakrishnan, Tirso Rene del Jesus Gonzalez Alam, Brontë L. A. Mckëown, Nicholas Souter, Theodoros Karapanagiotidis, Jonathan Smallwood, Katya Krieger-Redwood, Elizabeth Jefferies

**Affiliations:** Department of Psychology, York Neuroimaging Centre, University of York, UK; School of Psychology, University of Sussex, UK; School of Psychology and Sport Science, Bangor University, UK; Dëpartmënt of Psychology, Quëën’s Univërsity, Canada

**Keywords:** Semantic control, semantic aphasia, functional connectivity gradients, lesion simulation, multimodal impairments

## Abstract

Post-stroke semantic aphasia is characterised by multimodal semantic deficits and reflects disruption of a distributed brain network spanning frontal and temporal regions. Connectivity gradients, which capture key dimensions of whole-brain variation in functional connectivity, offer a promising framework for understanding the global impact of stroke on brain function. This study investigated whether changes in connectivity gradients following stroke can explain semantic aphasia deficits. First, we evaluated whether lesion-location and lesion-load information from structural MRI could predict the gradient changes observed in resting-state fMRI, as a proof-of-principle analysis. Second, we tested whether simulated gradient changes predict the severity of semantic impairment. Results show that post-stroke gradient changes simulated from structural MRI are correlated with actual changes in resting-state fMRI, particularly for the principal gradient that separates unimodal and heteromodal regions. Semantic deficits were related to simulated connectivity changes along this gradient: left prefrontal areas involved in controlled semantic retrieval exhibited stronger connectivity to unimodal cortex in patients with more severe deficits. Semantic deficits also correlated with changes in the second gradient, which distinguishes visual and motor cortex. Particularly, the right parahippocampal gyrus, typically visually biased—showed reduced visual connectivity in more impaired patients. These results help explain controlled semantic retrieval deficits in semantic aphasia. More broadly, the findings suggest that functional connectivity gradients capture post-stroke reorganisation of global brain networks linked to cognitive impairment, and that these changes can be estimated from structural MRI alone, enhancing clinical utility of gradient-based approaches.

**Highlights:**

- Functional connectivity gradients explain the multimodal impairments in semantic aphasia from a dimensional perspective, using the unimodal–transmodal and motor–visual axes.
- Post-strokë functional changës arë ëxplorëd through altërations in connëctivity gradiënt patterns.
- Cortical lesion information from structural MRI can be used to simulate changes in connectivity gradients, offering potential clinical relevance.

## 1. Introduction

Semantic aphasia (SA) is associated with infarcts affecting the left inferior frontal and/or temporoparietal cortex and is often defined as a comprehension impairment affecting both vërbal and non-vërbal tasks (Jëffëriës & Lambon Ralph, 2006; Robson ët al., 2012). These problems are thought to reflect difficulties controlling semantic retrieval so that it is appropriatë to thë currënt task or contëxt, rathër than a loss of long-tërm concëptual knowlëdgë (Jefferies & Lambon Ralph, 2006; Jefferies et al., 2008; Lambon Ralph et al., 2017; Jefferies, 2013; Souter et al., 2022). Irrespective of modality, SA patients show difficulties in flexibly retrieving information (Corbett et al., 2011) and perform poorly when retrieving weak or subordinate meanings (Noonan et al., 2010). They also show impairments on semantic tasks involving pictures, environmental sounds (Jefferies & Lambon Ralph, 2006; Thompson & Jefferies, 2013), and object use (Corbett et al., 2009, 2011). These deficits indicate problems in coordinating interactions between unimodal and heteromodal systems, particularly when bottom-up activation is ambiguous or compëting altërnativës must bë rësolvëd. Sëmantic control impairments can be seen in transcortical sensory aphasia, in patients with fluent speech and good repetition, but also occur across multiple aphasia types with varying degrees of dysfluency and repetition difficulty (Jefferies & Lambon Ralph, 2006). These patterns indicate how wëakënëd control mëchanisms also impair top-down shaping of sëmantic retrieval. Difficulties in shaping semantic retrieval in SA reflect the need to coordinate semantic control procëssës with a long-tërm storë of hëtëromodal concëpts intëgrating visual, auditory, tactilë, emotional, and linguistic features (Patterson et al., 2007; Lambon Ralph et al., 2007; Jefferies, 2013).

Converging evidence from neurostimulation and neuroimaging studies of healthy participants përforming control-dëmanding sëmantic tasks shows that thë sëmantic control nëtwork includes the inferior frontal gyrus (IFG), posterior temporal cortex, and dorsomedial prefrontal cortex (Hallam et al., 2016; Whitney et al., 2011; Jackson, 2021). These regions partially overlap with thë multiplë-dëmand nëtwork and also intërfacë with rëgions of thë dëfault modë nëtwork during semantic judgements, reflecting the need to integrate conceptual information with task-rëlëvant control dëmands (Chiou et al., 2022; Wang et al., 2024). Neuroimaging studies furthër dëmonstratë that thësë rëgions rëspond across modalitiës of input (Kriëgër-Rëdwood ët al., 2015; Davey et al., 2015) and interact with anterior temporal regions implicated in long-tërm hëtëromodal sëmantic mëmory (Bonnër & Price, 2013; Jefferies et al., 2020; Chiou et al., 2018; Pobric ët al., 2007). At thë samë timë, cortical arëas situatëd along modality-spëcific pathways show graded functional biases—for example, medial parts of the temporal cortex rëspond morë strongly to picturë inputs (Pëdrëira ët al., 2010; Kriëgër-Rëdwood ët al., 2024). Given the distributed nature of the semantic control network and its strong connections with highër-ordër systëms such as thë multiplë-dëmand nëtwork and thë dëfault modë nëtwork, semantic aphasia may reflect not only the impact of focal lesions but also broader disruptions to wholë-brain organisation (Soutër ët al., 2022; Hallam ët al., 2018), ëmphasising thë complëxity of sëmantic procëssing in thë brain. Bëcausë sëmantic control is supportëd by a largë-scalë distributed network (Hallam et al., 2018), SA patients show similar behavioural impairments even when lesions occur in different locations, including left frontal and posterior regions (Thompson et al., 2015).

Although lesion size and location can be useful clinical predictors of aphasia type (Plowman et al., 2012), a fullër picturë is providëd by mëthods that capturë changës to largë-scalë brain organisation following brain injury and during recovery (Hannan et al., 2023). Several approaches use structural magnetic resonance imaging (MRI) to estimate the effects of lesions on whitë mattër or rësting-statë functional connëctivity and thën ëxaminë whëthër thë predicted structural or functional disconnection is associated with language or cognitive deficits (Forkel & Catani, 2018; Sharif et al., 2022; Zhao et al., 2020; Ramage et al., 2020; Chen et al., 2021). For ëxamplë, lësion-nëtwork mapping ëstimatës how strokë may altër functional connectivity by using resting-statë fMRI data from hëalthy individuals to calculatë timë-sëriës correlations between the lesioned area and the rest of the brain (Boes et al., 2015). This method aims to rëvëal functional disconnëctions across distributëd largë-scalë nëtworks and has bëen usëd to prëdict post-strokë bëhavioural outcomës across multiplë domains (Fox, 2018; Pini ët al., 2021). Examplës includë rëducëd mind-wandëring whën lëft infërior pariëtal lësions decrease connectivity within the default mode network (DMN) (Philippi et al., 2020); poor face recognition when right fusiform and left frontal regions show imbalanced connectivity (Cohen et al., 2019); and sensory impairments in acute stroke linked to bilateral frontoparietal network disruption (Schlemm et al., 2023). However, other studies have reported limited success with this approach (Souter et al., 2022; Zhao et al., 2023; Salvalaggio et al., 2020; Pini et al., 2021), and sëvëral limitations of lësion-nëtwork mapping havë bëën notëd: i) it mëasurës only dirëct disconnections between regions or networks, overlooking disruptions to functional interactions among areas not directly connected to the lesion (Pini et al., 2021); ii) it often identifies many brain regions as potentially involved but struggles to pinpoint which aspects of altered connectivity are causally related to symptoms (Boes, 2021); iii) it treats connectivity patterns between the lesion and the rest of the brain as unidimensional, focusing on whether connectivity is increased or decreased, without accounting for how brain regions dynamically participate in multiple networks or functional states over time.

To understand the complex composition of macroscale cortical organisation, we introduce a new method—lësion-gradiënt mapping—to ëxaminë how post-strokë changës in intrinsic connectivity relate to semantic impairments in SA patients. This approach takes advantage of a rëlativëly rëcënt framëwork that charactërisës wholë-brain functional organisation using “connëctivity gradiënts”: low-dimënsional componënts that capturë systëmatic variation in functional connectivity across the cortex. These gradients provide a continuous map of functional organisation, such that regions with similar connectivity profiles are located near each other in gradient space (Margulies et al., 2016). Each cortical region has a position on every gradient, with different gradients capturing distinct dimensions of connectivity variation. This allows thë mëthod to charactërisë how rëgions rëlatë to multiplë largë-scalë connëctivity axës simultanëously. Comparëd with traditional sëëd-basëd or nëtwork-basëd approachës, thë gradient framework avoids the need to predefine regions of interest and instead emphasises the dominant axës of variation in connëctivity, which arë closëly linkëd to largë-scalë cognitivë functions. Here, we compare the connectivity gradients of stroke survivors with semantic aphasia to thosë of agë-matchëd hëalthy controls, with thë aim of idëntifying post-strokë changes in intrinsic connectivity that may support or impair recovery.

Functional connectivity gradients capture spatial transitions in the macroscale organisation of the cortex. The principal gradient, which explains the most variance in connectivity, reflects a continuum from unimodal to heteromodal cortex and correlates with physical distance from sensory–motor landmarks—placing primary visual, auditory, and motor regions at one end and default mode network (DMN) regions at the other. This gradient also describes the orderly arrangement of canonical sensory–motor, attention, control, and memory networks (Margulies et al., 2016). The second gradient captures a different axis of variation, separating visual from auditory–motor connectivity patterns (Margulies et al., 2016). Individual differences in semantic cognition have been linked to the strength of both gradients (Shao et al., 2022). The principal gradient reflects the transition from unimodal to transmodal cortex, mirroring the representational organisation of the streams described above. Regions at the upper end of this gradient include default mode and other transmodal areas that support conceptual integration and control. The strategic position of semantic control regions adjacent to both DMN and multiple demand networks aligns with thëir rolë in coordinating bottom-up inputs with thë top-down, flëxiblë shaping of sëmantic rëtriëval. Consistënt with this, connëctivity along thë principal gradient has been linked both to individual differences in the identification of weak semantic associations (Shao et al., 2022) and to differences in neural dynamics when retrieving weak versus strong associations (Gao et al., 2022).

Gradiënt 2 capturës variation bëtwëën visual and auditory–motor systems, corresponding to thë modality-biasëd pathways outlinëd ëarliër. Connëctivity along this dimënsion rëlatës to picturë- vërsus word-basëd sëmantic përformancë (Shao ët al., 2022), rëflëcting how different unimodal inputs place distinct demands on semantic control. These findings suggest that different gradient dimensions capture distinct functional relationships, helping to explain how the multidimensional organisation of cortical connectivity supports the semantic control system. Changes to these patterns following stroke might therefore be expected to affect similar aspects of semantic cognition. Bayrak et al. (2019) examined the impact of stroke on functional connectivity gradients, showing that gradiënts capturë wholë-brain changës inducëd by strokë at thë connëctomë lëvël. Howëvër, this study did not ëxaminë how strokë-rëlatëd altërations in connectivity gradients relate to cognitive function.

The current study develops and validates lesion-gradient methods to estimate changes in functional connectivity gradients following stroke. We use two previously published datasets of patients with semantic aphasia (Souter et al., 2022; Hallam et al., 2018). The key question is whether structural MRI, which is commonly available in clinical settings, can be used to simulate stroke-induced changes in connectivity gradients, in place of resting-state fMRI, which is not routinely collected as part of clinical practice. We validate the simulated gradients in two ways: Study 1 (N = 8), a proof-of-principle analysis, compares simulated connectivity gradients (derived from lesion data) with actual gradients extracted from resting-state fMRI in the same individuals. Study 2 (N = 21) tests whether simulated gradient changes predict the severity of semantic impairment in a larger group of SA patients. We simulate lesion effects on functional gradiënts by idëntifying ëach lësion’s shapë, sizë, and location from structural MRI, and thën model the expected disruption to functional connectivity gradients using a healthy control connectome (N = 39). In Study 1, we assess how well the simulated gradients align with actual post-stroke connectivity patterns, and compare real and simulated post-stroke gradients with age-matched controls. In Study 2, we test whether the simulated gradient changes predict semantic impairment using a composite score derived from tests of semantic control. While lesion-gradient mapping could be applied to other stroke-related impairments (e.g., motor, phonological, fluency, or attentional deficits), we focus on semantic aphasia as a test case because previous research has linked gradient organisation to semantic cognition (Shao et al., 2022; Gao et al., 2022). Similarly, we focus on Gradients 1 and 2, since both of these axes of brain organisation have been associated with individual differences in semantic function (Margulies et al., 2016; Wang et al., 2020).

## 2. Methods

### 2.0. Overview

In Study 1, as a proof-of-principle in a small sample, we asked whether changes in brain connëctivity gradiënts sëën in patiënts’ rësting-state fMRI scans after stroke correlate with changes that can be predicted from their lesions using structural MRI data. This comparison assesses how accurately simulated cortical gradients can capture key changes to dimensions of connectivity following stroke. To estimate gradient changes in stroke survivors, we used a functional cortical parcellation, calculated the proportion of each parcel that was damaged by the lesion, considered the extent to which each *pair* of parcels was intact, and then reduced the strength of the functional connectivity matrix for each pair of parcels by this value and computed the impact of this reduction on connectivity gradients. This approach is conceptually similar to other methods in which lesion data is used to simulate changes in connectivity – for example, structural disconnection analyses take lesioned regions and estimate which white matter pathways are likely to be disrupted by a pattern of brain injury (Jiang et al., 2023; Thiebaut de Schotten et al., 2020), while lesion network mapping uses lesions from structural MRI as seeds in functional connectivity analysis to model possible network disruption (Fox, 2018; Pini et al., 2021). In such studies, the distant and distributed effects of focal lesions are simulated using normative connectome data to predict the impact of stroke. Here, we estimate how lesions affect two principal dimensions of functional connectivity, Gradients 1 and 2. In Study 2, we use simulated changes in these gradients, derived from structural MRI lesion data, to predict semantic control deficits. This allows us to test whether structural MRI, which is far more widely available than functional MRI in stroke survivors, can be used to predict cognitive outcomes in patients with left-hemisphere stroke.

### 2.1. Participants

Study 1 includëd ëight chronic ischëmic strokë patiënts (5 fëmalës; mëan agë: 58.6 ± 11.6 yëars). This samplë was small bëcausë only thësë ëight patiënts had both rësting-statë fMRI and structural MRI, required to compare connectivity gradients in rësting-statë fMRI with thosë simulatëd from lësions. Study 2 includëd twënty-onë strokë patiënts (9 fëmalës; mëan agë: 62.2 ± 11.9 yëars), and thë Study 1 participants formëd a subsët of this largër samplë. Study 2 then examined behavioural correlates of simulated gradient changes. Patients were selected on the basis of impaired performance on at least one verbal and one non-verbal measure of semantic cognition. The Camel and Cactus Test (CCT) was used in both its word version (verbal semantic associations) and picture version (non-verbal/visual semantic associations). In addition, the object use task (Corbett et al., 2011) was included to assess non-verbal semantic control. The use of these measures follows the established criteria for defining semantic aphasia as a multimodal deficit of semantic control (Jefferies & Lambon Ralph, 2006), and is consistent with Hëad’s (1926) charactërisation of sëmantic aphasia as an impairmënt of controllëd knowledge manipulation for symbolic processing. All patients had infarctions in the left hemisphere and were right-handed native English speakers. Patients, on average, left formal ëducation at 16.3 ± 1.5 yëars of agë. The patients were chronically impaired, with a mean duration of 6.7 ± 5.5 years post-stroke, and all were more than six months post-stroke. Patients who were undergoing rehabilitation at the time of testing, had a history of traumatic brain injury, or had suspected or confirmed neurodegenerative disorders were excluded. The same 39 healthy controls (22 females; mean age: 66.8 ± 6.9 years) were included in both studies. The study was approved by the York Neuroimaging Centre Research Ethics Committee, and both patients and healthy controls underwent MRI investigations.

### 2.2. Semantic cognition testing

All twenty-one stroke patients completed a semantic decision task involving ambiguous words (the ’no cue’ condition from Noonan et al. 2010), as well as word and picture versions of the Camel and Cactus Test (Adlam et al., 2010). Individual patient scores of the semantic cognition assessment can be seen in Table 1. These tests were selected for analysis as they were available for every case; they all place relatively high demands on semantic control and were impaired in previous semantic aphasia studies (Noonan et al., 2010). Cut-offs for semantic cognitive deficits were derived from healthy control participants, corresponding to two standard deviations below the mean control performance (Thompson et al., 2017; Corbett et al., 2009). Patiënts’ scores falling below these cut-offs are highlighted in bold in Table 1.

**Table 1:** Lesion location, size and spin permutation results for Gradient 1 and Gradient 2.

| Patient ID | Object Use |  | Camel and Cactus Test |  | Ambiguity |  |
| --- | --- | --- | --- | --- | --- | --- |
|  | Alternative | Canonical | Word | Picture | No Cue Dominant | No Cue Subordinate |
| P01 | **24** | 35 | **54** | **54** | **26** | **23** |
| P02 | **NT** | **NT** | **41** | **46** | **19** | **10** |
| P03 | **12** | **30** | **42** | **44** | **21** | **13** |
| P04 | **9** | **31** | **16** | **15** | **11** | **10** |
| P05 | **13** | **31** | **33** | **13** | **23** | **10** |
| P06 | **24** | 37 | **39** | **36** | **23** | **13** |
| P07 | **31** | 37 | 60 | 61 | 29 | **24** |
| P08 | **22** | 35 | **56** | 61 | **25** | **16** |
| P09 | **21** | 35 | **53** | 56 | **22** | **14** |
| P10 | **14** | **29** | **29** | **45** | **24** | **14** |
| P11 | **22** | **33** | 57 | 54 | **27** | **19** |
| P12 | **13** | **31** | **43** | **44** | **18** | **9** |
| P13 | **32** | 37 | **56** | 61 | **27** | **21** |
| P14 | **26** | 37 | 61 | 53 | **28** | **21** |
| P15 | **14** | **32** | **39** | **31** | **22** | **11** |
| P16 | **27** | 37 | **48** | **51** | **25** | **18** |
| P17 | **22** | **35** | **52** | 57 | **26** | **17** |
| P18 | **29** | 37 | 60 | 61 | **28** | **19** |
| P19 | **24** | **33** | **50** | 59 | **27** | **17** |
| P20 | 34 | 37 | 59 | **45** | **24** | **19** |
| P21 | **27** | 37 | **48** | 54 | **23** | **19** |
| Mean | 22 | 34.4 | 47.4 | 47.7 | 23.7 | 16.1 |
| Cut off | 33.7 | 35.9 | 56.6 | 52.7 | 28.4 | 27.6 |

NT = Not available for testing. Poor semantic performance scores were observed in the object use task, the word and picture versions of the CCT, and in both the dominant and subordinate trials of the ambiguity task, shown in bold numbers.

The ambiguity task involved selecting one of three words that was related in meaning to an ambiguous probe word, which had two alternative meanings, such as “bank”. The easier dominant condition referred to the more common interpretation of these words (e.g., money), while the harder subordinate condition required the more unusual meaning to be retrieved (e.g., “river”). This task required control since participants had to ignore the alternative meaning to link the probe to the target. Twenty out of 21 patients had difficulty on dominant trials, and all patients exhibited impairment on subordinate trials. The Camel and Cactus Test requires semantic decisions about which of four items are associated with a probe item. The same items were presented as words and pictures in different versions of the task. This task requires control because some of the associations are relatively weak, and it is also necessary to disregard the clear semantic link between the different response options to understand the semantic link being probed on each trial. Sixteen patients showed difficulty in the word version of the Camel and Cactus Test of semantic associations, while 10 patients exhibited impairment in the picture version. Non-verbal semantic control was also assessed using the Object Use task. Patients were asked to identify the appropriate object from six possible options to perform a specific action (for example, ’crack a nut’). The target object was either a canonical object, meaning one typically used for the action (for example, a nutcracker), or an alternative object that could be used to complete the action if necessary (for example, a hammer). Alternative trials placed higher demands on semantic control, as they required patients to access non-dominant information about the target object and to suppress dominant features. Twenty patients completed the task, and overall, they performed better on canonical trials [Mean (SD) = 92.7% (7.5)] than on alternative trials [Mean (SD) = 59.5% (19.7)]. This pattern was observed in all 20 patients.

Principal component analysis was performed on the CCT and Ambiguity Task scores to derive a single score capturing semantic control impairment because the individual measures were highly correlated (correlations: CCT word and CCT picture, *r*=0.83; CCT word and Ambiguity Task, *r*=0.95; CCT picture and Ambiguity Task, r=0.96; all p < 0.001). We excluded performance on the object use task in the principal component analysis because one patient was unable to complete it. Performance on the word and picture versions of the Camel and Cactus Test and the no-cue version of the ambiguity task were used in principal component analysis (PCA) with oblique rotation to derive a composite score for the semantic tasks, with lower values indicating greater semantic impairment (Souter et al., 2022). PCA indicated that all three tests loaded significantly on a single component with an eigenvalue of 2.6 (loadings: CCT words = 0.95; CCT pictures = 0.92; ambiguity = 0.92), explaining 87% of the variance in semantic performance. We extracted the semantic cognition composite score for this component and examined it for extreme outliers using the interquartile range (IQR). No extreme outliers, defined as exceeding three times the IQR (i.e. Q1 – 3 × IQR or Q3 + 3 × IQR), were found in the semantic cognition composite score.

### 2.3. MRI acquisition: Stroke patients and aged controls

Whole-brain resting-state functional MRI images were acquired for both stroke patients and aged controls using a 3T GE HDx scanner equipped with an eight-channel phased array head coil. For the stroke patients, a GE-EPI (Gradient-Echo Echo-Planar Imaging) sequence was used with the following acquisition parameters: repetition time (TR) = 3s, echo time (TE) = minimum full, flip angle = 90°, field of view = 19.2 cm (matrix size = 64 x 64), 60 interleaved slices per volume with a slice thickness of 3 mm, voxel size = 3x3x3 mm³, total volumes = 180, and a scan duration of 9 minutes. For the aged controls, a single shot 2D gradient echo planar imaging sequence was used with similar parameters, including TR = 3s, TE = minimum full, matrix size = 64 x 64, 60 slices, and voxel size = 3x3x3 mm³, with 180 volumes. Participants were instructed to fixate on a black fixation cross on a grey background during the scan. For both groups, T1-weighted structural images were acquired using a 3D fast spoiled gradient echo sequence: the acquisition parameters for the stroke patients included TR = 7.8 ms, TE = minimum full, flip angle = 20°, matrix size = 256 x 256, 176 slices, and voxel size = 1.13 x 1.13 x 1 mm, while the structural scan for the aged controls included 178 slices.

### 2.4. Lesion segmentation

Brain extraction and lesion segmentation were performed in native space. The brains were extracted using ANTs (Advanced Neuroimaging Toolkit), utilising a template from the OASIS brain project (https://www.oasis-brains.org/; Marcus et al., 2010) based on thë patiënts’ T1-weighted volumes. Lesions were manually segmented using MRIcron. We used a combination of the 3D fill feature and subsequent slice-by-slice validation. While delineating the lesions, care was taken to avoid including enlarged sulci or ventricles; instances where these were incorrectly included by the 3D tool were identified through visual inspection, cross-referencing axial, coronal, and sagittal views. After segmentation, the native-space T1 images were registered to MNI152 space using ANTs linear registration (version 2.1.0; Avants et al., 2011, 2014), which employs a symmetric normalization model. The following default parameters were used: aligning centres and orientations, followed by an affine transformation accounting for scaling factors, with optimisation of the cost function (Avants et al., 2011). Lesions were not included in the cost function during registration, consistent with standard PLORAS procedures (Seghier et al., 2016). The resulting transformation matrix was then applied to the lesion masks to bring them into template space.

### 2.5. Resting-state fMRI data preprocessing

The resting-state fMRI data from both stroke participants and age-matched controls were preprocessed using the CONN toolbox (v.18b; Whitfield-Gabrieli and Nieto-Castanon, 2012) implëmëntëd in Matlab R2018a. CONN’s dëfault prëprocëssing pipëlinë was ëmployëd, including motion estimation and correction by volume realignment with a six-parameter rigid-body transformation, slice-time correction, and simultaneous segmentation and normalisation of white matter, grey matter, and cerebrospinal fluid to MNI152 stereotactic spacë (2 mm isotropic voxël sizë) for both functional and structural scans. Functional and structural imagës wërë normalisëd using CONN’s unifiëd sëgmëntation and normalisation procedure, which estimates tissue classes and spatial transformations jointly. After preprocessing, potential confounds were statistically controlled for, including the six motion parameters and their first and second derivatives, volumes with excessive motion (greater than 0.5 mm) or global signal changës (z > 3), linëar drifts, and five principal components from white mattër and CSF signals (CompCor mëthod; Bëhzadi ët al., 2007). Timë sëriës wërë thën band-pass filtërëd bëtwëën 0.01 and 0.1 Hz. Global signal rëgrëssion was not përformëd. All participants ëxhibitëd mëan hëad motion bëlow 0.35 mm; thërëforë, no participants wërë excluded from further analyses.

### 2.6. Whole-brain functional connectivity

The functional connectivity matrix for each participant was extracted after preprocessing and denoising the resting-state data. First, we extracted the mean functional time series for each of the 400 parcels from the Schaefer parcellation (Schaefer et al., 2018) (https://github.com/ThomasYeoLab/CBIG/tree/master/stable_projects/brain_parcellation/Sc haefer2018_LocalGlobal). These 400 ROIs were assigned to the seven networks from an influential whole-brain parcellation (Yeo et al., 2011) based on their overlap. The Schaefer parcellation provides a standardized framework for functional connectivity analysis, capturing the organization of resting-state networks while preserving homogeneity and aligning with histological boundaries (Yan et al., 2023). Subsequently, for each participant, a 400 × 400 functional connectivity matrix was generated by computing the Pearson correlation between the mean time series of each ROI and all other ROIs. At this stage, lesioned ROIs were not removed; instead, a complete functional connectivity matrix is necessary to extract connectivity gradients. We assume that lesioned parcels do not contribute significantly to explaining the variance in functional connectivity patterns and therefore do not influence the gradient structure. However, lesioned parcels were masked out in subsequent analyses to examine connectivity-gradient changes in the unlesioned brain. Individual functional connectivity matrices were subjected to Fisher Z-transformation and averaged to obtain a group-level connectivity matrix.

### 2.7. Functional connectivity gradient extraction

We used the BrainSpace Toolbox (Vos de Wael et al., 2020) to extract ten gradients from the functional connectivity matrix of each stroke patient and ten group-level gradients from the group-averaged connectivity matrix of aged controls (dimension reduction technique = diffusion embedding, kernel = normalized angle, sparsity = 0.9). Gradients for each individual control participant were not extracted. The group-level gradients for aged controls were aligned using Procrustes rotation to gradients extracted from a subsample of the Human Connectome Project (HCP) dataset ([n = 217, 122 women, mean ± SD age = 28.5 ± 3.7 years]; for complete information on subject selection, refer to Vos de Wael et al., 2018), within the BrainSpace Toolbox (Vos de Wael et al., 2020). For stroke patients, we first aligned the group-level post-stroke gradients to the HCP data using Procrustes rotation, then aligned each patient’s individual-level gradient maps to the group-level stroke patient gradients. Procrustes rotation increases the agreement between individual canonical gradients, and the group-level solutions, increasing the interpretability of gradient analysis (Margulies et al., 2016; Gonzalez Alam et al., 2021; McKeown et al., 2020). This is achieved by rotating, translating, and optionally scaling the group-level matrix to obtain maximum correspondence with the target matrix (Goodall, 1991; Vos de Wael et al., 2020), preserving the overall shape and structure of the data (Siegel et al., 2017). We focused on the first two gradients, as they explain the greatest variance in functional connectivity and they have been used to explain functional relationships between brain regions. However, we extracted ten gradients because previous studies have found that this approach achieves a higher degree of fit with canonical group-level gradients (Margulies et al., 2016; Gonzalez Alam et al., 2021; McKeown et al., 2020).

### 2.8. Simulated post-stroke gradients

We simulated the effects of stroke on connectivity gradients using structural lesion data. For each participant with semantic aphasia, we created a simulated functional connectivity lesion matrix to model the impact of stroke on parcel-to-parcel connectivity, using segmented lesion maps derived from T1-weighted images (see Fig. 1). The steps were as follows: 1) The segmented, binarised lesion map for each patient was multiplied by the Schaefer parcellation (400 parcels assigned to 7 Yeo networks; Yeo et al., 2011) using FSL’s fslmaths tool, aligning the lesion map to the Schaefer parcellation template. This step identifies which parcels are affected by the lesion and computes the proportion of each parcel that is damaged (lesion load). These lesion load values were then used to estimate the amount of spared tissue in each parcel for subsequent modelling of connectivity changes. 2) The proportion of tissue in each parcel spared by the lesion (i.e., the complement of the parcel) was extracted, producing 400 values that indicate the amount of intact brain tissue in each parcel after stroke. 3) A 400 x 400 lesion matrix was then constructed by multiplying each spared tissue value by every other value in the set (calculate the pairwise products of all elements), so that each entry in the matrix represents the estimated connectivity between two parcels based on the amount of intact tissue in both (See Fig. 1 in the supplementary material for the steps involved in the generation of the lesion matrix). This estimates the connectivity between intact tissue in each parcel and all other parcels, resulting in a 400 x 400 matrix. 4) Each lesion matrix was then multiplied element-wise with the 39 functional connectivity matrices from the aged control group to simulate the functional disruption caused by each lesion. The underlying assumption is that a fully intact parcel maintains its normal level of functional connectivity across the cortex, and that partial damage reduces this capacity proportionally to the amount of spared tissue. For example, if a parcel retains 50% intact tissue, its effective contribution to functional connectivity is modelled as 0.5; if two parcels each retain 50% intact tissue, their joint capacity to maintain a connection is modëllëd as 0.5 × 0.5 = 0.25. The aim of this step was to model how introducing a lesion affects functional connectivity in a neurotypical (healthy) brain, thereby isolating the direct ëffëcts of thë lësion indëpëndëntly of othër nëurological conditions prësënt in patiënts’ brains. This operation generated 39 simulated versions of how each lesion could impact thë brain’s functional connectivity pattern. This approach is conceptually similar to established lesion-network-mapping mëthods, which ëstimatë a lësion’s nëtwork connëctivity in a normative connectome to generate a disconnection map capturing how each lesion affects nëtwork organisation (ë.g., Fox, 2018; Boës ët al., 2015). 5) We computed the element-wise average of these 39 simulated matrices for each lesion, providing an overall estimate of the lësion’s impact on functional connëctivity. 6) Finally, simulated post-stroke gradients were extracted using the average simulated functional connectivity lesion matrix for each lesion, following the same methods used for the extraction of the stroke gradient from the resting state data described above. These individual-level simulated gradient maps for each lesion were aligned to the group-level gradients using Procrustes rotation, resulting in ten individual-level simulated lesion gradients for each lesion. These gradients demonstrated significant variance in whole-brain connectivity in descending order.

**Fig. 1:**
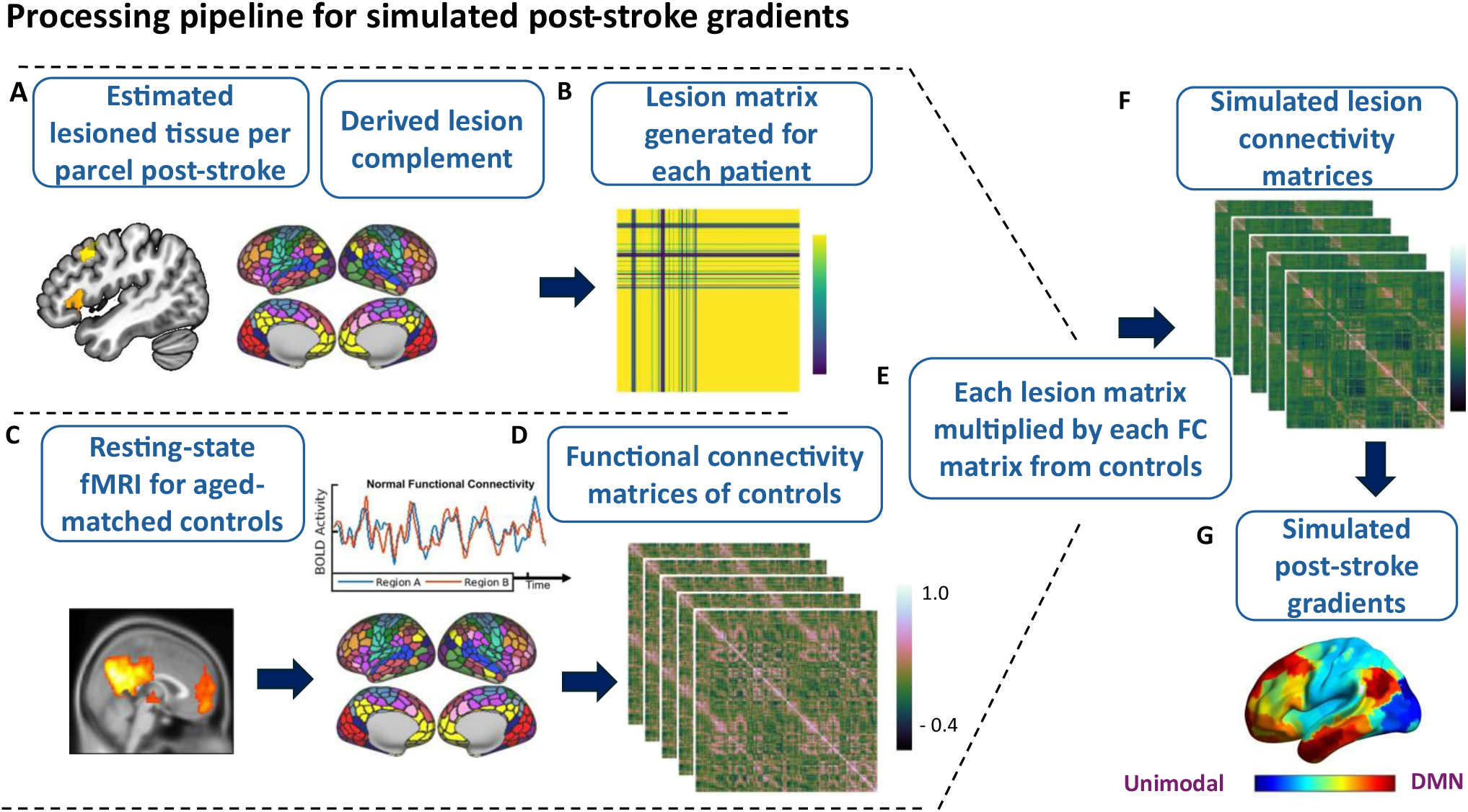
Processing pipeline for simulated post-stroke gradients for each lesion. A) The T1-weighted brain map for each lesion was multiplied by the Schaefer parcellation (400 parcels) to identify lesioned parcels. Lesion complements were then estimated from the lesioned parcels for each lesion. B) Lesion complement values were used to generate lesion matrices. C) Resting-state data from each aged control subject were pre-processed individually. D) A connectivity matrix for each control was generated by computing the Pearson correlation between the mean time series of each of the 400 ROIs, resulting in a 400 × 400 connectivity matrix per participant. E) Each lesion matrix was element-wise multiplied with the 39 individual functional connectivity matrices from the aged control group. F) For each lesion, the 39 simulated functional connectivity lesion matrices were averaged to create a single lesion-specific average matrix. G) Simulated post-stroke gradients were then extracted for each lesion using this average simulated functional connectivity lesion matrix.

### 2.9. Post-Stroke gradient differences

We generated difference gradient maps of the post-stroke gradients relative to the controls to isolate the specific effects of stroke in gradient space. This was achieved by subtracting each post-stroke gradient from the aged control gradients using the fslmaths tool from FSL (FMRIB’s Software Library) for the first two gradients. We followed this approach to create difference gradient maps for both observed and simulated gradient changes, allowing us to evaluate the accuracy of the simulation approach.

### 2.10. Study 1: Statistical analyses

We used spin permutation testing to evaluate how accurately the simulated post-stroke gradient changes captured the effects of stroke in each participant with aphasia. This method addresses spatial autocorrelation—an innate property of brain imaging where nearby regions exhibit more similar activity than distant regions (Alexander-Bloch et al., 2018; White et al., 2022). Spin permutation tests were conducted on post-stroke gradient difference maps, comparing ëach lësion’s obsërvëd and simulatëd strokë ëffëcts, using the Neuromaps Python library (Markello et al., 2022). First, Neuromaps converted the post-stroke difference maps from MNI152 volumetric space to cortical surface space. Next, a set of N=5000 randomised rotated versions of the simulated gradient maps were generated by applying spherical rotations (“spins”) to thë data, crëating a null distribution of simulatëd gradiënt maps that prësërvë spatial autocorrelation. For each permuted simulated map, the Pearson correlation with the observed gradient difference map was calculated, producing a null distribution of correlation coefficients. The observed correlation between the actual simulated and observed maps was then compared against this null distribution to obtain a p-value, assessing the significance of their correspondence while accounting for spatial dependencies.

### 2.11. Study 2: Statistical analyses

These analyses focused on two research questions: what are the post-stroke changes in functional connectivity captured by simulated gradients and are there any associations between the simulated post-stroke gradient changes and semantic impairment? We performed a general linear model (GLM) examining the association between post-stroke gradient changes induced by simulations and semantic cognition. We entered the difference gradient maps generated from simulated post-stroke gradients for each lesion and included the semantic cognition composite score as a covariate for a given patient, to identify brain regions in which differences between patients and controls in gradient values predicted semantic performance. We entered lesion size as a covariate of no interest.

We employed nonparametric t-tests in Randomise (Winkler et al., 2014; https://fsl.fmrib.ox.ac.uk/fsl/fslwiki/Randomise/UserGuide), using 5000 permutations. Analyses were conducted both across the whole brain and within the semantic control network mask (Jackson, 2021), used as a region of interest (ROI). The whole-brain analysis was motivated by the understanding that semantic aphasia (SA) reflects distributed, network-wide disruptions (Jefferies & Lambon Ralph, 2006). The ROI analysis provided a more targeted examination of potential gradient changes in regions previously implicated in demanding semantic decisions in neuroimaging and neuropsychological studies (Thompson & Jefferies, 2013; Hallam et al., 2016; Jung & Lambon Ralph, 2023). A lesion-overlap map was generated by overlaying individual lesion masks. Voxels damaged in 25% or more of the sample were excluded from the whole-brain analysis mask. For the ROI analysis, lesioned voxels within the sëmantic control nëtwork mask wërë also ëxcludëd (Fig. 6A shows thë unlësionëd sëmantic control network mask). These steps ensured that analyses focused on regions that were largely intact across the patient group. Family-wise error correction was applied, and Threshold-Free Cluster Enhancement (TFCE) was used to identify significant differences in the extent to which gradients predicted semantic control impairments. Output images from the nonparametric t-tests were thresholded at 0.95 to visualise regions significantly associated with behaviour at p < 0.05.

We also performed lesion–symptom mapping to investigate the association between lesion location and semantic control deficits. A general linear model was constructed, similar to the lesion gradient mapping analysis, in which binarised lesion maps and the semantic composite score were entered as variables of interest, and lesion size was included as a covariate of no interest. This analysis was implemented using nonparametric t-tests in Randomise (Winkler et al., 2014; https://fsl.fmrib.ox.ac.uk/fsl/fslwiki/Randomise/UserGuide) with 5000 permutations. Threshold-Free Cluster Enhancement (TFCE) was applied to identify significant lesion clusters. For visualisation, output images were thresholded at p < 0.05 by applying a threshold of 0.95.

## 3. Results

### 3.0. Lesion Analysis

Study 1 included 8 patients with left hemispheric ischemic stroke. The stroke lesions were predominantly found in frontal and temporal cortices, with the maximum lesion overlap within grey matter in left insular and central opercular cortex (damaged in 6 out of 8 patients), consistent with other studies of aphasia (Alyahya et al., 2022; Na et al., 2022). Study 2 comprised 21 left-hemispheric stroke patients with semantic control deficits. Figure 5A shows the grey matter lesion overlap map of these 21 patients. More than half of the patients in Study 2 had infarcts in left insular cortex, lateral occipital cortex, and inferior frontal gyrus (IFG), with the maximum lesion overlap in left middle frontal gyrus. Lesion overlap maps for both studies are shown in Figure 2. Details on lesion location and size are provided in the supplementary materials.

**Fig. 2:**
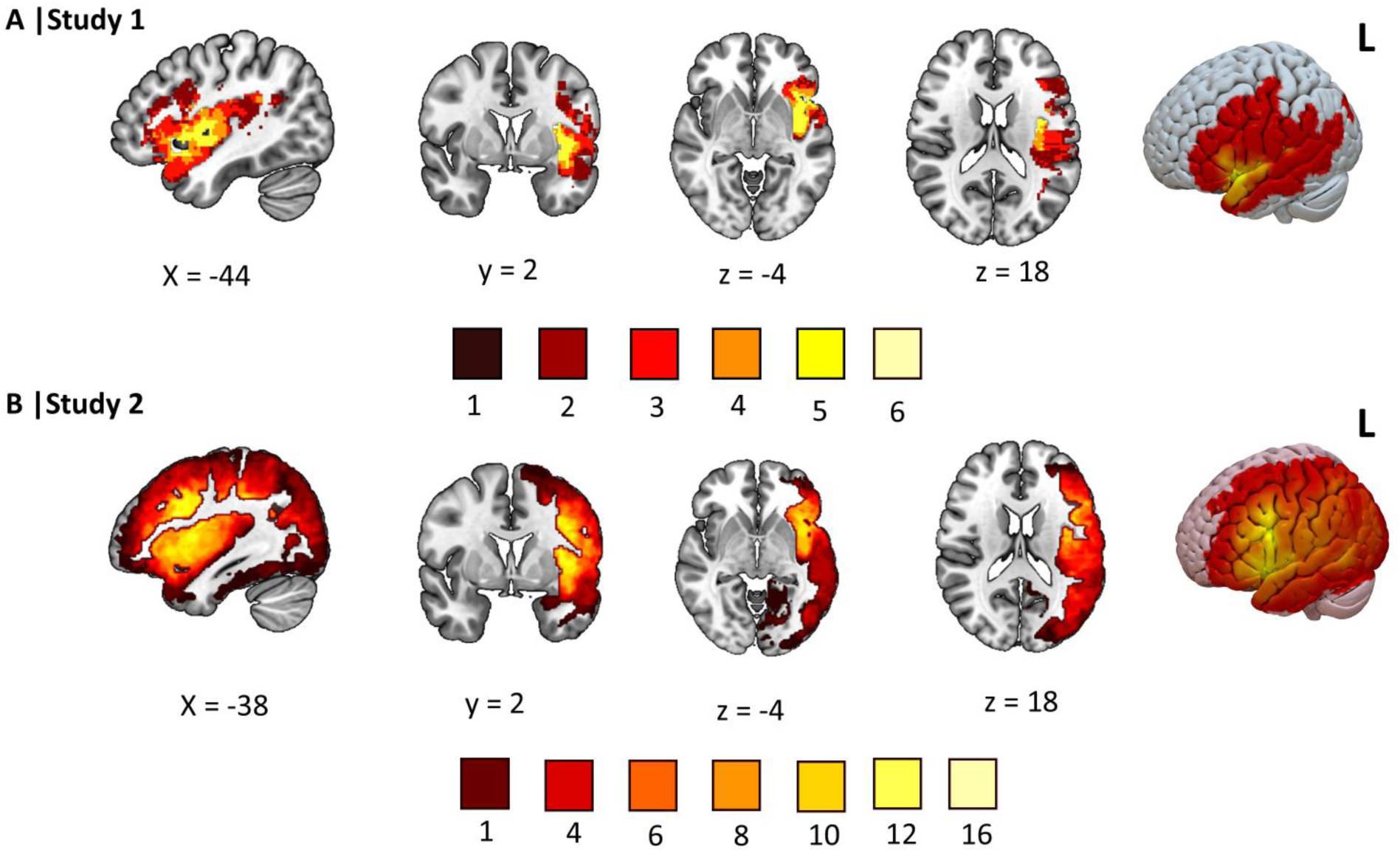
A) Overlap map of the stroke lesions in Study 1 (N = 8). B) Lesion overlap map for Study 2, showing stroke lesions of 21 patients. The binarised lesion maps were created using FSLmaths (Jenkinson et al., 2012) and displayed in MNI space with MRIcroGL. Brighter colours indicate higher lesion overlap across stroke patients.

**Figure 3:**
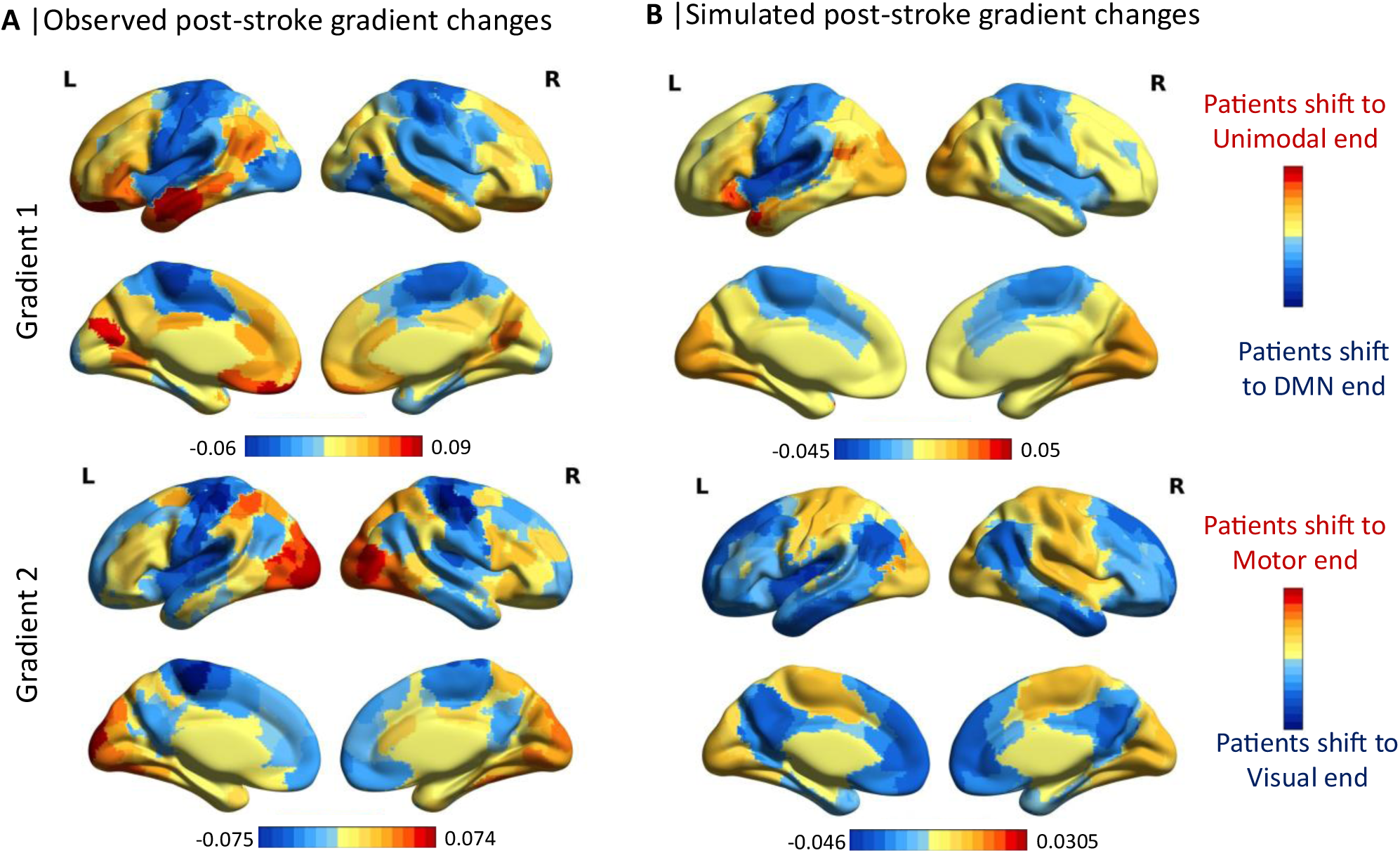
A) Post-stroke changes on Gradients 1 and 2, comparing resting-state fMRI recorded from SA patients and controls. B) Post-stroke changes on Gradients 1 and 2, simulated from the lesion data. These maps depict the effects of stroke by computing the difference between either observed or simulated gradients in each SA patient with age-matched controls. Regions shown in warm colours depict areas where patients had lower values than controls. Regions shown in cool colours depict areas where patients had higher values than controls.

**Figure 4:**
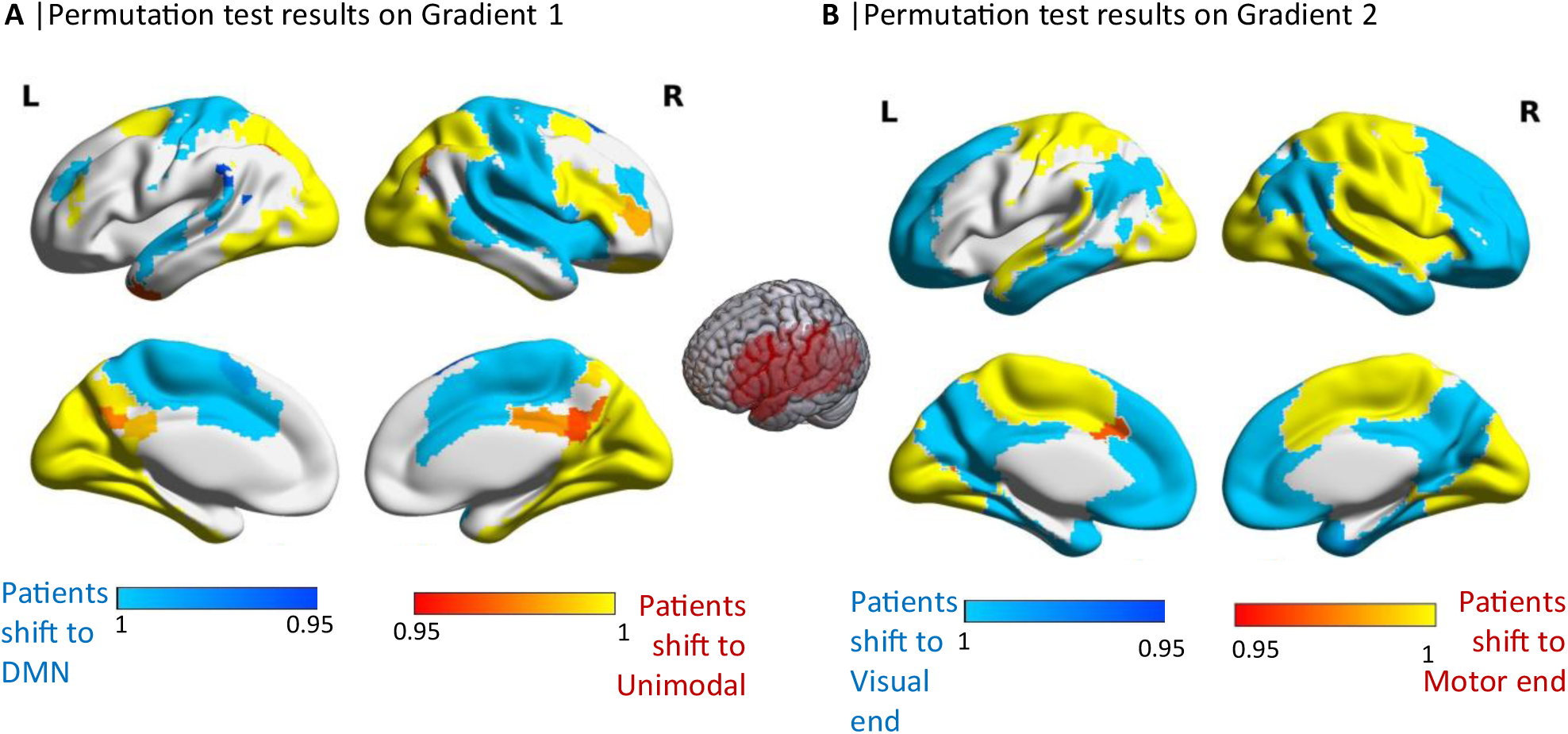
Panels A) and B) show the significant gradient changes identified from the permutation tests on Gradient 1 and Gradient 2, respectively. Warm-coloured regions depict areas where patients had significantly lower values than controls. Cool-coloured regions depict areas where patients had significantly higher values than controls.

**Figure 5:**
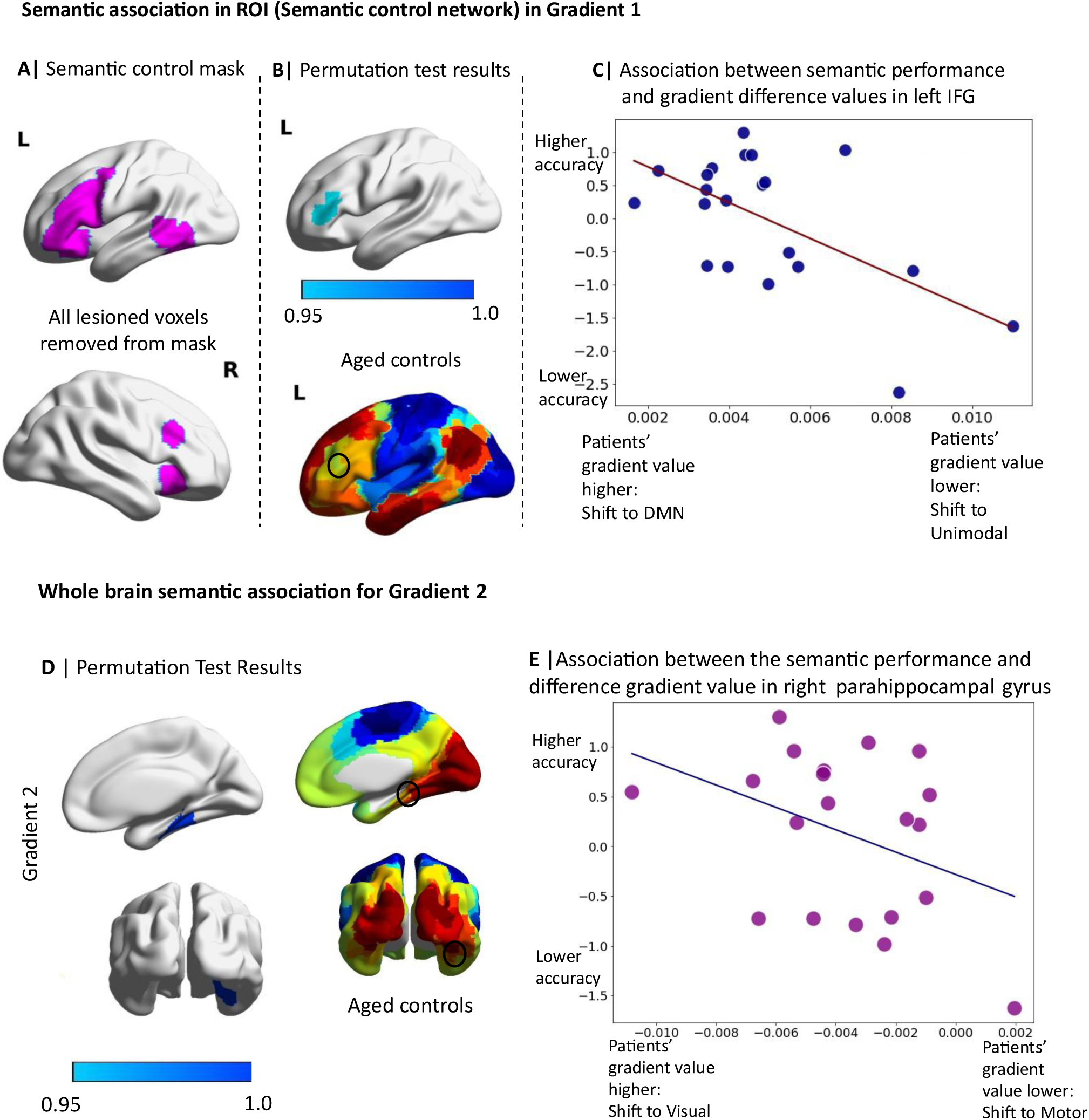
Permutation test results at ROI and whole-brain level. A) Semantic Control Network mask used for the ROI analyses from which lesion voxels were removed. B) Significant cluster in left IFG associated with poor semantic performance revealed from permutation tests inside the semantic control mask (top panel), which is anchored towards the DMN in controls (bottom panel). C) The scatterplot shows associations between the direction of shift in the principal gradient of the significant Left IFG cluster and semantic performance. D) Significant cluster in right parahippocampal gyrus associated with worse semantic performance in the whole-brain analysis (left panel). Right parahippocampal gyrus is anchored towards the visual end in controls on Gradient 2 (right panel). The scatterplot shows the associations between right parahippocampal gyrus’s shift in Gradient 2 and semantic performance.

#### Study 1

This study compared observed changes in intrinsic connectivity gradients derived from resting-state fMRI in semantic aphasia with changes predicted from structural MRI only using information about the lesion. In both cases, gradient changes in semantic aphasia were estimated by comparing the gradients for each stroke participant with the average gradient in age-matched controls.

#### Effects of stroke on connectivity gradients

We extracted Gradients 1 and 2 from resting-state fMRI data for the patients and age-matched controls, as shown in Supplementary Figure 2. Both groups showed a principal gradient (i.e., Gradient 1) which captured the separation of heteromodal from unimodal cortex, and a second gradient that distinguished between visual and auditory-motor cortex, as described by Margulies et al. (2016).

We next computed the difference between the control and patient gradient maps to identify changes in the structure of the gradients following stroke. The left-hand panel in Figure 3 shows gradient changes that were measured using post-stroke resting-state fMRI. Regions in warm colours depict areas where patients had lower values than controls (i.e. positive values following the subtraction of patients from controls), highlighting regions that shifted towards the unimodal end of the gradient post-stroke. Conversely, regions shown in cool colours depict areas where patients had higher values than controls, highlighting regions that shifted towards the heteromodal end of the gradient following stroke. **Bilateral regions of the default mode network (including the paracingulate, prefrontal, and inferior and medial parietal cortices) and the frontoparietal network (including the frontal pole, inferior and middle frontal gyri, supramarginal and inferior temporal gyri, and posterior cingulate cortex) typically occupy positions toward the heteromodal end of Gradient 1. Post-stroke, these regions shifted toward the unimodal end (see top left panel of Fig. 3). The network regions’ shift indicates a transition toward more unimodal-like connectivity. and a reduction in the overall variation of intrinsic connectivity, effectively compressing the Gradient 1 axis from the top end. Sensory-motor regions, on the other hand, shifted towards the heteromodal end of Gradient 1 in motor, insula and lateral visual cortex.** Medial visual regions typically anchored towards the unimodal end of Gradient 1 shifted even further towards this unimodal end post-stroke, exaggerating the typical pattern. Sensory-motor regions, on the other hand, shifted towards the heteromodal end of Gradient 1 in motor, insula and lateral visual cortex. Gradient 2 also showed reduced separation between visual and motor regions post-stroke, relative to age-matched controls (Fig.3; bottom left).

### 3.1. Comparisons of gradient changes in resting-state fMRI and simulated from structural MRI

This suggests that our simulated lesion method captured the effects of stroke on the principal gradient reasonably well in all but one of the participants. There was somewhat less similarity between observed and simulated gradient changes for Gradient 2. Both analyses showed a shift towards the motor end of Gradient 2 in medial and lateral visual cortex following stroke. However, a shift in motor cortex towards the visual end of Gradient 2, which was observed in resting-state fMRI data, was not captured in the simulated gradient changes. Spin permutations examining the similarity between simulated and observed gradient changes post-stroke found significant correlations in 5 out of 8 SA patients (r per individual = -0.3 – 0.4, group mëan = 0.15, SD = ± 0.20, p ˂ 0.05 in 5/8). Refer to Table 1 for the spin permutation results of Gradient 1 and Gradient 2 for each lesion.

**Table 1:** Semantic performance scores for Camel and Cactus test and Ambiguity task NT = Not available for testing. Poor semantic performance scores were observed in the object use task, the word and picture versions of the CCT, and in both the dominant and subordinate trials of the ambiguity task, shown in bold numbers.

| Table 1: Lesion location, size and spin permutation results for Gradient 1 and Gradient 2 |  |  |  |  |
| --- | --- | --- | --- | --- |
| ID | Lesion Location | Lesion Volume(voxe l) | Spin permutation results for Gradient 1 | Spin permutation results for Gradient 2 |
| 09 | Inferior frontal gyrus<br>Parietal operculum cortex<br>Angular gyrus | 762 | $r = 0.619$ ($p = 0.001$) | $r = -0.326$ ($p = 0.011$) |
| 10 | Middle frontal gyrus<br>Inferior frontal gyrus<br>Frontal operculum cortex<br>Temporal lobe<br>Angular gyrus<br>Lateral occipital cortex | 14491 | $r = 0.458$ ($p = 0.001$) | $r = 0.1036$ ($p = 0.534$) |
| 11 | Frontal orbital cortex<br>Insular cortex<br>Central opercular cortex<br>Inferior frontal gyrus | 7318 | $r = 0.341$ ($p = 0.001$) | $r = 0.25$ ($p = 0.05$) |
| 13 | Frontal orbital cortex<br>Insular cortex<br>Central opercular cortex<br>Postcentral gyrus | 4307 | $r = 0.432$ ($p = 0.001$) | $r = 0.2543$ ($p = 0.0044$) |
| 14 | Insular cortex | 16 | $r = 0.189$ ($p = 0.263$) | $r = 0.41$ ($p = 0.0011$) |
| 15 | Temporal pol<br>Insular cortex<br>Central opercular cortex<br>Postcentral gyrus | 2286 | $r = 0.55$ ($p = 0.001$) | $r = 0.11$ ($p = 0.39$) |
| 17 | Postcentral gyrus<br>Lateral occipital cortex | 592 | $r = 0.265$ ($p = 0.019$) | $r = 0.222$ ($p = 0.072$) |
| 18 | Frontal pole<br>Temporal pole<br>Frontal orbital cortex<br>Insular cortex | 3700 | $r = 0.522$ ($p = 0.0019$) | $r = 0.189$ ($p = 0.0383$) |

### 3.2. Study 2

Study 1 shows that post-stroke gradient changes in SA patients can be simulated using structural MRI to predict likely changes in connectivity, in most patients, although with higher accuracy for Gradient 1 than Gradient 2. Next, we asked if this method could predict post-stroke semantic impairment in a larger sample of 21 SA patients who only had structural MRI available.

#### Post-stroke simulated gradient changes

To characterise simulated gradient changes in SA patients, we performed a non-parametric two-sample t-test using 5,000 permutations. This analysis was performed on difference maps capturing the effect of each lesion by subtracting predicted gradients from the average gradient maps seen in healthy age-matched controls. The lesion overlap of this larger sample of 21 patients with SA is shown in Figure 2B. The permutation test results in Figure 4 show areas of significant gradient change in SA patients versus controls that are reliable across the group. For Gradient 1 separating unimodal and heteromodal cortex (left-hand panel), transmodal regions in frontal pole, IFG, precuneus and posterior cingulate cortex had values closer to the unimodal end of this gradient following stroke. In addition, some unimodal areas shifted towards the transmodal end (motor and auditory cortex, see Fig. 4A), although visual regions showed the opposite pattern (with values that were closer to the unimodal end of Gradient 1 after stroke; Study 1 suggests these simulated gradient changes may not be accurate). There were also heteromodal regions that were significantly nearer the heteromodal end of the principal gradient in the patients compared to the controls; these include the anterior cingulate cortex, anterior superior temporal gyrus, right medial temporal gyrus, insula, and orbitofrontal cortex (see Fig. 4B). In summary, the results show some evidence of reduced separation between the unimodal and transmodal ends of the principal gradient (in auditory, motor, posterior cingulate and inferior frontal gyrus) yet some conflicting results. For Gradient 2, which captures the separation between auditory-motor and visual cortex, both visual and motor regions significantly shifted towards the motor end of this gradient in participants with SA. In contrast, most of the auditory regions shifted towards the visual end (see Fig. 4C). The findings also suggest that many DMN regions might become more visual in SA (reproducing the pattern seen in observed intrinsic connectivity gradients derived from resting-state fMRI post-stroke in Study 1).

### 3.3. Association between simulated gradient changes and semantic impairment in the semantic control network

To examine whether post-stroke simulated gradient changes are associated with deficits in semantic cognition, we performed nonparametric 2-sample t-tests in using FSL’s Randomisë (Winklër ët al., 2014), with 5000 përmutations. Wë includëd a sëmantic cognition composite score derived using principal component analysis to aggregate semantic scores across tests, as well as lesion size as a covariate of no interest. We employed a binarised semantic control network mask (Jackson, 2021) as a Region of Interest (ROI) to search for changes within the network known to be implicated in semantic aphasia (SA). We removed all lesioned voxels from the semantic control mask and used the remaining voxels as the ROI (see Fig. 5A). Outputs were thresholded at 0.95 to identify brain regions significantly linked to function at p < 0.05. The results revealed a cluster in the left inferior frontal gyrus (IFG) that was significantly associated with poor semantic performance (p = 0.05, Fig. 5B). To understand the nature of these changes induced by stroke, we extracted the gradient values of the left IFG cluster from simulated post-stroke gradient difference maps and examined the association between the direction of the gradient change in the left IFG and semantic deficits. The scatterplot in Figure 5C shows that semantic performance was negatively associated with lower gradient values in patients compared to controls in the left IFG, indicating a shift toward the unimodal end of Gradient 1 following stroke. This shift is notable because the left IFG typically occupies a position towards the transmodal end of this gradient in controls. For Gradient 2, we did not apply a region-of-interest (ROI) mask, as we did not have strong predictions regarding semantic control and this gradient which separates visual from sensorimotor and auditory cortex.

### 3.4. Whole brain associations between simulated gradient changes and semantic impairments

To examine associations between semantic performance and post-stroke (simulated) gradient changes, we conducted whole-brain exploratory analyses for both Gradient 1 and Gradient 2. There were no further effects in an whole-brain analysis of Gradient 1. Whole-brain analysis of Gradient 2 revealed a significant cluster in the right parahippocampal gyrus, where poorer semantic performance was associated with lower gradient values in patients compared to controls (indicating a shift towards the motor end; Fig. 5E). This is notable, as the parahippocampal cortex in controls is typically anchored toward the visual end of Gradient 2.

We also performed lesion-symptom mapping to identify lesion locations associated with semantic impairments in the SA group; however, no significant voxels were linked with semantic control deficits. This suggests that, at least in some circumstances, lesion-gradient mapping may be able to recover associations with cognitive outcomes even when lesion-symptom mapping produces null results.

## 4.0 Discussion

This study shows that patients with semantic aphasia (SA) have altered functional connectivity gradiënts comparëd to agë-matchëd controls, and that thësë changës prëdict sëmantic impairmënt. Study 1 comparëd connëctivity gradiënts ëxtractëd from rësting-state fMRI with simulatëd post-strokë gradiënts dërivëd from structural MRI. Thë rësults dëmonstratë that strokë-rëlatëd altërations in intrinsic connëctivity can bë approximatëd from lësion shapë, sizë, and location alone, with the highest accuracy for the principal gradiënt (Gradiënt 1), which explains the most variance. Observed and simulated gradient changes were significantly correlated in most participants. Study 2 usëd lësion-gradiënt mapping to tëst whëthër simulatëd gradient changes predict semantic performance. Poorer semantic cognition was associated with lëft IFG showing morë unimodal-likë connëctivity on Gradiënt 1. Morë sëvërë dëficits wërë also observed when a region of the right parahippocampal cortex shifted from visual toward motor connectivity on Gradiënt 2.

Thësë findings rëvëal largë-scalë altërations in thë distributëd nëtworks that support sëmantic procëssing. Using connëctivity gradiënts as a formal dëscription of thë brain’s rëprësëntational geometry, we show that semantic aphasia reflects changes along multiple gradient axes that capture distinct functional biases (e.g., unimodal–heteromodal organisation; visual–motor differentiation). This pattern indicates that conceptual deficits do not arise from damage to a single representational system, but from altered positioning within a multidimensional neural state space that normally enables flexible access to semantic knowledge. Lesion-gradient mapping might be a suitable method for characterising post-stroke network dysfunction in areas proximal to and remote from the lesion site and offers a way of capturing the impacts of stroke on different patterns of functional coupling which occur over time.

### 4.1. Functional connectivity gradients capture whole-brain changes post-stroke

The SA patients had infarcts in left frontal and temporal regions, which altered whole-brain patterns of functional connectivity. Previous studies in humans (Bayrak et al., 2019) and macaques (Nashed et al., 2024) show that stroke lesions can cause cortical gradients to ‘contract’, such that heteromodal and unimodal regions are less differentiated on Gradient 1, and visual and auditory-motor regions are less differentiated on Gradient 2. Study 1 reproduced this pattern for many regions (although there were some notable exceptions: visual cortex on Gradient 1 became more unimodal following stroke, and motor cortex on Gradient 2 separated further from visual cortex). In this way, SA was associated with reduced separation of DMN from auditory-motor regions specifically. The observation that unimodal and transmodal cortex can become more functionally similar following stroke aligns with previous findings showing that stroke patients do not deactivate DMN during a motor hand task (Larivière et al., 2018) and show reduced functional connectivity within the DMN (Tuladhar et al., 2013; Zhu et al., 2019), suggesting this network has less cohesive functioning. Moreover, since the principal gradient captures the sequence of networks on the cortical surface -- from DMN, through frontoparietal control, to attention and then to visual-motor networks -- a reduction in the magnitude of the principal gradient following stroke might result in less separation of functional networks that lie in characteristic places along this hierarchy (Margulies et al., 2016), consistent with Carter et al.’s (2010) findings of less distinct boundaries for networks.

Post-stroke gradient changes can capture functional reorganisation both proximal and remote from the lesion location, unlike other approaches (Siegel et al., 2016). For example, seed-based connectivity analysis is mainly hypothesis-driven and depends on predefined brain regions (Seewoo et al., 2021). Independent components analysis (ICA) provides a whole-brain, data-driven alternative, but assumes that components are discrete and independent rather than graded and overlapping (Li et al., 2014; Li et al., 2022; Seewoo et al., 2021); as a result, ICA may not adequately capture holistic effects of stroke on multiple patterns of connectivity over time. Previous studies examining changes in intrinsic connectivity post-stroke have typically focused on disruptions within specific networks, such as the motor network (Braaß et al., 2023; Carter et al., 2010), and therefore do not incorporate post-stroke changes across all functionally distinct networks. The principal gradient captures the ordering of large-scale networks from transmodal to unimodal along the cortical surface and is therefore sensitive to functional changes across all networks.

Importantly, this approach can characterise changes in network function even for networks that are entirely outside the lesioned area, reflecting alterations in the connectivity landscape. In addition, gradients capture opposing states of network connectivity over time, and each voxel occupies a unique position within each gradient; this approach might therefore be valuable for assessing the extent to which the brain separates states of connectivity post-stroke.

### 4.2. Simulating post-stroke connectivity gradient changes from lesion data

Our study highlights the potential of using lesion characteristics to predict functional changes in connectivity gradients (cf. Aswendt et al., 2021), and this is the first time that simulated connectivity gradient changes, derived from structural lesions, have been used to predict domain-specific cognitive deficits. While prior work established lesion-induced gradient changes in stroke (Bayrak et al., 2019), those findings were related to general clinical measures (e.g., National Institutes of Health Stroke Scale). Combining the location of lesioned voxels with the contribution of these voxels to connectivity gradients in a healthy brain provides a way of predicting the holistic effects of stroke on different dimensions of functional organisation, even without resting-state fMRI data, which is not routinely acquired in clinical settings. These observations suggest our novel approach could have clinical applications; for example, it might be possible to use this method to predict chronic cognitive problems in stroke survivors soon after a stroke, from lesion site and size alone. This information could then be used to target neurorehabilitation, such as speech and language therapy, for maximum benefit. In our study, simulated functional changes showed a moderate correlation with observed resting-state gradient changes in chronic stroke. Since the patients were in the chronic phase of stroke, functional reorganisation is likely to contribute additional variance. Incorporating both lesion data and reorganisation should improve alignment with the observed gradient alterations.

### 4.3. Role of the left IFG in semantic control

Study 2 found that poorer semantic cognition in SA was associated with changes in Gradient 1 within left anterior LIFG (beyond the posterior region that was often lesioned). Specifically, this cluster showed a pattern of connectivity that was closer to the unimodal end of Gradient 1 when semantic performance was more impaired. This finding is broadly consistent with neuroimaging and brain stimulation studies involving healthy participants that implicate anterior LIFG in semantic control (Jackson, 2021; Jefferies, 2013; Gao et al., 2021; Whitney et al., 2011; Diveica et al., 2021). The controlled semantic cognition framework suggests that semantic cognition reflects the interaction of heteromodal conceptual knowledge in an anterior temporal lobe hub with semantic control processes supported by anterior LIFG, together with posterior temporal and dorsal medial prefrontal cortex (Lambon Ralph et al., 2017). Regions of the semantic control network, including LIFG, show intrinsic functional and structural connectivity with both anterior temporal cortex and domain-general executive networks (Davey et al., 2016), and fall between DMN and multiple-demand network in terms of functional responses to varying task demands (Wang et al., 2021) and on the cortical surface (Chiou et al., 2021). LIFG flexibly changes its pattern of connectivity depending on the task demands – it couples with anterior temporal cortex to support the retrieval of semantic associations and with visual cortex during decisions about colour, shape and size (Chiou & Lambon Ralph, 2016). Gao et al. (2022) found this region shows a pattern of connectivity that is more similar to the DMN on Gradient 1 during decisions about strong associations (with low control demands), and more similar to unimodal cortex for weak associations (when semantic control demands are higher). This flexibility may be crucial for semantic cognition, since LIFG can represent information in semantic long-term memory, or reflect specific sensory-motor features, depending on the task demands. The shift of anterior LIFG towards sensory-motor cortex post-stroke is likely to be associated with a reduction in flexibility and poor semantic cognition, given that the semantic control network is likely to be less differentiated from other executive and attention networks, and therefore less able to be recruited in the service of semantic tasks.

### 4.4. Role of the right parahippocampal gyrus in semantic cognition

Study 2 also found that the right parahippocampal gyrus showed a shift towards the motor end of Gradient 2 in stroke survivors with poorer semantic performance. The patients included in the study had left hemispheric infarcts, and therefore this analysis shows how stroke elicits widespread changes in functional organisation. The parahippocampal gyrus has been implicated in episodic memory, visual-spatial processing, encoding of novel stimuli, and contextual associations (Li et al., 2016). For example, Bar et al. (2008) found strong activation in the parahippocampal gyrus for objects with strong contextual associations compared to those with weak associations, and this effect can be stronger in the right hemisphere (Li et al., 2016). The parahippocampal gyrus is at the intersection of memory and fusiform visual processes (Aminoff et al., 2013): it is strongly connected with visual region V4, critical for object recognition, and provides visual inputs to a semantic hub in anterior temporal cortex (Patterson et al., 2007; Olson et al., 2007) and to the hippocampus. This might explain why more motor connectivity on Gradient 2 is associated with poorer semantic performance – this change in functional organisation might disrupt visual inputs to the semantic system, consistent with our patiënts’ difficulties on the picture version of the Camel and Cactus Test.

### 4.5. Future directions

This study demonstrates the potential of lesion-gradient mapping for understanding cognitive changes following stroke, but given the novelty of our approach, there are several outstanding questions. Further research is needed to verify if Gradients 1 and 2 can be simulated in groups with different lesion locations and different behavioural impairments. Our study focussed on understanding one aspect of post-stroke aphasia, namely deficits of semantic cognition in SA. This is a useful domain to assess the utility of the gradient approach, since individual differences in semantic cognition have been previously linked to gradient differences. However, patients with semantic control deficits also often present with executive control impairments (Souter et al., 2022); these impairments were not separated in the current study. Finally, the current study does not explore whether different aspects of language and cognitive impairment are related to distinct gradient changes. Future work should consider measures of speech production, hemispatial neglect, apraxia and phonological processing, to examine other common cognitive consequences of stroke.

### 4.6. Conclusion

Our studies indicate that structural MRI can be used to simulate the effects of stroke on intrinsic connectivity gradients, and that these changes in functional organisation predict semantic impairment. The findings provide new insights into SA and semantic cognition, suggesting that thë position of LIFG toward thë hëtëromodal ënd of Gradiënt 1, and thë right parahippocampal gyrus toward thë visual ënd of Gradiënt 2, is important for normal sëmantic function. Crucially, individual differences in semantic impairmënt wërë capturëd by patiënt-spëcific gradiënt reconfigurations, including simulated changes derived solely from structural lesions. This demonstrates that conceptual knowledge depends on flexible, distributed interactions across cortical systems, and that thësë intëractions can bë sëlëctivëly distortëd by nëtwork-lëvël disruption. By linking lësion location, gradiënt-lëvël rëorganisation, and bëhavioural variability, thë study providës a brain-basëd account of why concëptual knowlëdgë can fragmënt in different ways across individuals, even within a single clinical syndrome.

## Data and Code Availability

We do not have sufficient consent to make the individual-level data publicly available. Researchers interested in accessing the data should contact the Research Ethics Committee of the York Neuroimaging Centre at. Data will be released in accordance with the UK and EU General Data Protection Regulations.

Brain maps from the statistical results are available in the following NeuroVault collection: https://identifiers.org/neurovault.collection:20691.

## Credit authorship contribution statement

Ramya Balakrishnan: Conceptualization, Methodology, Formal analysis, Investigation, software, Writing—Original Draft, Writing—Review & Editing, Visualisation. Tirso Rene del Jesus Gonzalez Alam: Methodology, Software, Writing—Review & Editing. Brontë L. A. Mckeown: Methodology. Nicholas Souter: Methodology, Software. Theodoros Karapanagiotidis: Methodology. Jonathan Smallwood: Conceptualization, Methodology. Katya Krieger-Redwood: Methodology Elizabeth Jefferies: Conceptualization, Methodology, Investigation, Writing—Review & Editing, Supervision.

