## Supplementary material for "Lesion-Gradient Mapping in Semantic Aphasia: Comparisons of Observed and Simulated Effects of Stroke on Connectivity Gradients"

### Supplementary Materials

### Methods

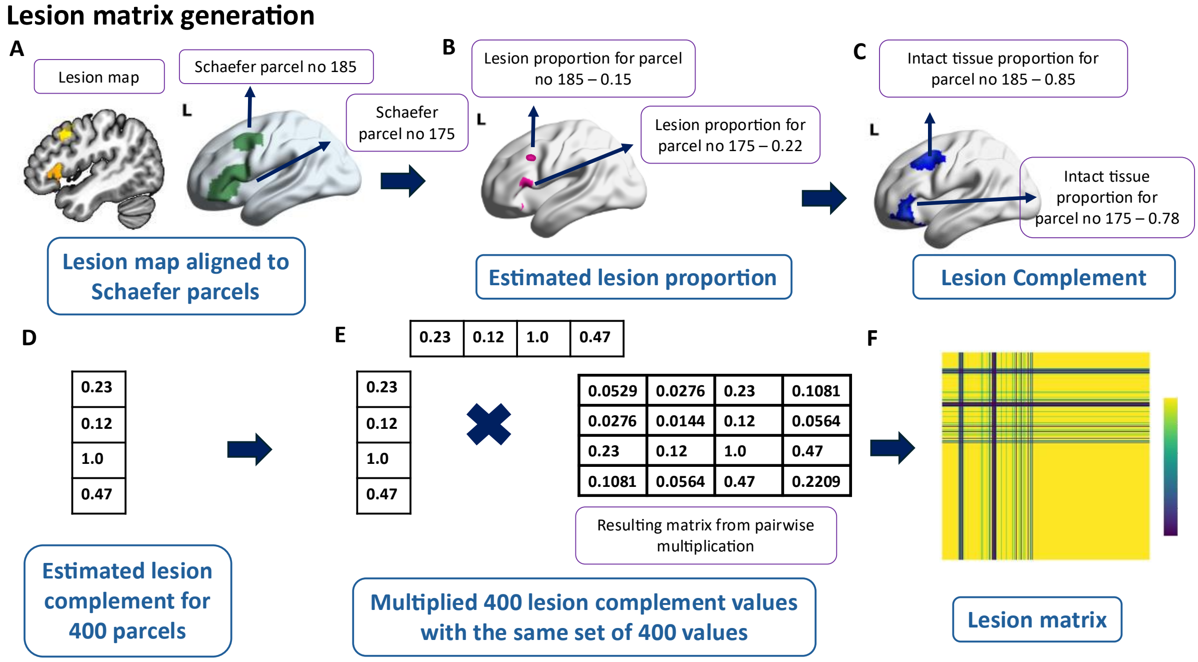


Fig.1 Steps followed for generating the lesion matrix: A) The lesion map was aligned to the Schaefer parcellation. B) The lesion proportion was estimated for each parcel. C) For each lesion, the lesion complement (spared tissue) per parcel was calculated from the lesion proportion. D) For each lesion, lesion complements were estimated for all 400 parcels. E) The 400 lesion complement values were multiplied by the same set of 400 values, resulting in a 400 x 400 lesion matrix (Example values from the matrix shown). F) Lesion matrix generated.

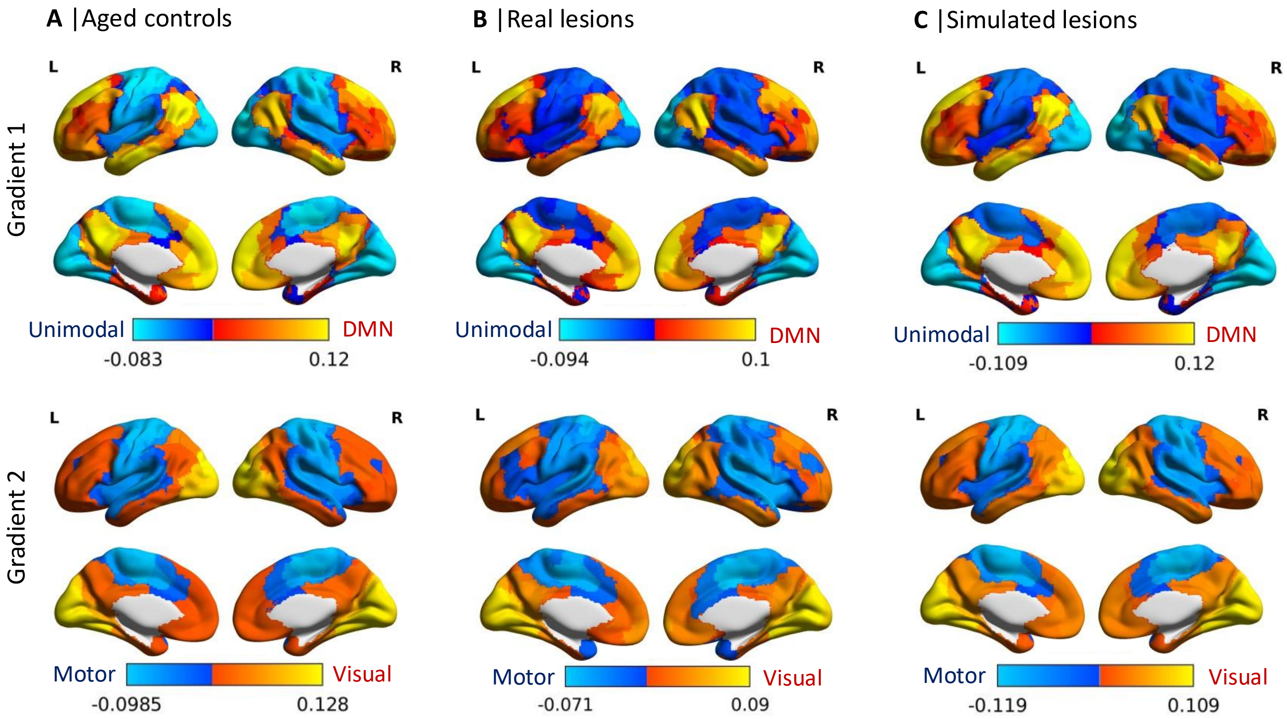


Fig.2: A) The topographic functional organisation of each gradient in aged controls. B) & C) The topographic cortical organisation of each functional gradient from resting state scans of stroke patients and simulated gradients from structural lesion data, respectively: On gradients 1 and 2, regions that occupy the bottom end of the gradient are shown in cool colours, and regions that are anchored towards the top end of the gradient are shown in warm colours.

Table 1explains the lesion location and size of the lesions. Lesion volume is reported in terms of the number of voxels (1 × 1 × 1 mm3).

**Table 2: Stoke infarct lesion location and size**

| **ID** | **Hemisphere** | **Lesion Location** | **Lesion Volume(voxel)** |
|---|---|---|---|
| 01 | Left | Lateral occipital cortex<br>Occipital cortex<br>Insular cortex | 4039 |
| 02 | Left | IFG<br>Lateral occipital cortex<br>Temporal fusiform cortex | 8766 |
| 03 | Left | Intracalcarine cortex<br>Parahippocampus | 168 |
| 04 | Left | Lateral occipital cortex<br>Motor cortex<br>Prefrontal cortex(lateral)<br>IFG | 13463 |
| 05 | Left | Lateral occipital cortex<br>Precuneous<br>Posterior cingulate cortex | 2596 |
| 06 | Left | Insular cortex<br>Motor cortex<br>Frontal pole<br>IFG | 6943 |
| 07 | Left | Lateral occipital cortex<br>Precuneous<br>Posterior cingulate cortex | 8317 |
| 08 | Left | Insular cortex<br>IFG<br>Middle frontal gyrus<br>Temporal cortex<br>Supramarginal gyrus | 13907 |
| 09 | Left | Middle frontal gyrus<br>Angular gyrus<br>Lateral occipital cortex | 762 |

| | | | |
|---|---|---|---|
| 10 | Left | Middle frontal gyrus<br>IFG<br>Insular cortex<br>Lateral occipital cortex<br>Postcentral gyrus<br>Temporal cortex | 14491 |
| 11 | Left | IFG<br>Temporal pole<br>Insular cortex<br>Supramarginal gyrus | 7318 |
| 12 | Left | IFG<br>Middle frontal gyrus<br>Frontal orbital cortex<br>Insular cortex<br>Temporal lobe | 7190 |
| 13 | Left | Frontal orbital cortex<br>Insular cortex<br>Post central gyrus | 4307 |
| 14 | Left | Insular cortex | 16 |
| 15 | Left | Insular cortex<br>Temporal pole | 2286 |
| 16 | Left | IFG<br>Lateral occipital cortex<br>Angular gyrus<br>Supramarginal gyrus<br>Postcentral gyrus<br>Temporal cortex | 8809 |
| 17 | Left | Middle frontal gyrus<br>Lateral occipital cortex | 592 |
| 18 | Left | IFG<br>Frontal orbital cortex<br>Insular cortex<br>Temporal pole | 3700 |
| 19 | Left | Superior frontal gyrus<br>Middle frontal gyrus | 16807 |

| | | | |
|---|---|---|---|
| | | IFG<br>Motor cortex<br>Insular cortex<br>Temporal pole<br>Supramarginal gyrus | |
| 20 | Left | Superior frontal gyrus<br>Middle frontal gyrus<br>IFG<br>Insular cortex<br>Motor cortex | 6590 |
| 21 | Left | Angular gyrus<br>Postcentral gyrus | 597 |
